# Horizontal gene transfer of a key translation protein has shaped the polyproline proteome

**DOI:** 10.1101/2023.10.19.563058

**Authors:** Tess E. Brewer, Andreas Wagner

## Abstract

Prolines take longer than other amino acids to be incorporated into nascent proteins and cause ribosomes to stall during translation. This phenomenon occurs in all domains of life and is exacerbated at polyproline motifs. Such stalling can be eased by elongation factor P (EFP) in bacteria. We discovered a potential connection between horizontally transferred EFP variants and genomic signs of EFP dysfunction. Horizontal transfer of the *efp* gene has occurred several times throughout the bacterial tree of life, and such transfer events are associated with the loss of otherwise highly conserved polyproline motifs. In this study, we pinpoint cases of horizontal EFP transfer among a diverse set of bacterial genomes and examine the consequences of these events on genome evolution in the phyla Thermotogota and Planctomycetes. In these phyla, horizontal EFP transfer is not only associated with the loss of conserved polyproline motifs, but also with the loss of entire polyproline motif containing proteins, whose expression is likely dependent on EFP. In particular, three proteases (Lon, ClpC, and FtsH) and three tRNA synthetases (ValS, IleS1, IleS2) appear highly sensitive to EFP transfer. The conserved polyproline motifs within these proteins all reside within, or in close proximity to ATP binding regions, some of which have been shown to be crucial to their function. Our work shows that the horizontal transfer of EFP has left genomic traces that persist to this day. It also implies that the process of ‘domesticating’ a horizontally transferred *efp* gene can perturb the overall function of EFP.

## Introduction

Proline-rich regions are important for protein function. They are over-represented in protein domains that are important for interactions with other proteins and with nucleic acids^1^. They are also associated with high cellular complexity^2,3^. However, proline-rich regions cause a problem during mRNA translation that is universal to all domains of life. Due to its uniquely rigid structure, proline is the slowest amino acid to be incorporated into proteins during translation. Proline causes translating ribosomes to pause or “stall” on mRNA, which slows down protein synthesis. This problem is compounded when an amino acid sequence contains multiple adjacent prolines (polyproline motifs, ≥2P). The severity of this stalling is determined by several factors. They include the identity of the surrounding amino acids (particularly the two amino acids before and one after the prolines e.g., XXPPX)^4^, the translation initiation rate^5^, and the position of the polyproline motif within a protein^6^. Polyproline motifs can have a dramatic impact on translational rate in highly expressed proteins, where they cause ribosomal queuing, wreak havoc on translational efficiency, and unlink the coupling between transcription and translation^5,7,8^.

To mitigate this problem, species in all three domains of life encode proteins that minimize the impact of ribosomal stalling at polyproline motifs^9^. In bacteria, this protein is called elongation factor P (EFP). EFP binds to the ribosome between the peptidyl-tRNA binding and tRNA-exiting sites. From here, EFP interacts with the peptidyl-transferase center, accelerating the formation of proline-proline bonds^9^. In many species, EFP must be post-translationally modified to function properly. While several different types of modification are known, these modifications all reside at the same position within EFP—on an amino acid positioned at the tip of a conserved loop region in domain I of the protein^9^. In most Gammaproteobacteria, the modified amino acid is a lysine (L34). It is modified by β-(*R*)-lysylation^10^ in chemical reactions that are catalyzed by the proteins EpmA and EpmB, as well as by a hydroxylation step of unknown function carried out by EpmC^11^. In some Firmicutes the modified amino acid is also a lysine (L32, equivalent to L34 in *E. coli*), but it is modified by the attachment of a 5-aminopentanol group carried out in several steps by the proteins YmfI, YnbB, and GsaB^12,13^. In Betaproteobacteria, the modified amino acid is an arginine (R32, equivalent to L34 in *E. coli*), which is rhamnosylated by the protein EarP^14^. Some bacteria, such as the Actinobacteria, use an unmodified EFP that can be identified by the presence of a conserved proline loop^15^. Lastly, some species contain an EFP paralog of unknown function known as YeiP^16^. YeiP is most often found in genomes encoding an EpmAB type EFP and is common among Gammaproteobacteria^14^.

Because EFP is important for efficient translation, loss of the *efp* gene can have dramatic phenotypic consequences. They include reduced growth rate^8,17–20^, loss of motility^21,22^, loss of virulence^17,18,20,22^, reduced antibiotic resistance^20,23^, and in the case of *Acinetobacter baumannii*^22^ and *Neisseria meningitidis*^24^, death. Many of these phenotypes are caused by the under-expression of proteins containing polyproline motifs^1^. In some proteins, altering such motifs to reduce the severity of ribosomal stalling can ease EFP loss-of-function phenotypes. For example, swarming motility can be restored to *Bacillus subtilis efp* mutants by modifying a ribosomal stalling polyproline motif in the flagellar C-ring component FliY^21^. In other proteins, polyproline motifs cannot be altered without substantial negative impacts on protein function. Examples include a polyproline motif within the glucose importer component EII^Glc^ in *Corynebacterium glutamicum*, which cannot be altered without inactivating the protein^25^ and a proline triplet in valine tRNA synthetase that is crucial for efficient and accurate tRNA charging in *E. coli*^26^. In other words, while some proteins can be modified to reduce their reliance on EFP, other proteins seem unavoidably dependent on EFP for normal expression.

In this study, we investigate a genomic mystery. In the course of a previous analysis^3^, we discovered that some species within the bacterial phylum Planctomycetota do not encode a proline triplet in valine tRNA synthase that is crucial to the enzyme’s function and was thought to be universally conserved across all domains of life^26^. Where the valine tRNA synthases of all other known forms of life encode a ‘PPP’ motif^26^, these species instead encode ‘PLP’. Pulling at this thread, we found that loss of this motif coincides with the appearance of a horizontally transferred EFP variant in these Planctomycete genomes. Relevant in this regard is that modifying polyproline motifs to reduce the severity of the ribosomal stall they cause during protein synthesis can restore normal expression in EFP deletion mutants^21^. Taken together, these observations suggest a connection between the horizontal transfer of EFP and global EFP dysfunction. EFP transfer has been observed previously^14,16^, but never studied in detail. For example, the EarP type EFP is believed to have originated within the Betaproteobacteria, but is also sporadically present within some Gammaproteobacteria, Thermotogota, Planctomycetes, Spirochaetes, and Fusobacteria^14^. In the underlying transfer events, the proteins that post-translationally modify EFP are generally co-transferred together with the EFP-coding gene in a single operon. This supports previous speculations that different EFP ‘types’ co-evolve with their modification systems^14,16^.

In this study, we examined how frequently horizontal transfer of EFP occurs within bacterial genomes. We find that horizontally transferred EFP does not often coexist with the ‘native’ ortholog, that is, the variant of EFP that existed within the recipient genome before the transfer. We also examined in more detail the genomes of species within two phyla that encode horizontally transferred EFP, the Planctomycetota and Thermotogota. In these species, horizontal transfer of EFP is consistently linked with loss of otherwise highly conserved polyproline motifs, such as the nearly universally conserved proline triplet in valine tRNA synthetase. Our work shows that the horizontal transfer of EFP is associated with proteome remodeling to alter polyproline motifs. In some species, this leads to the loss of entire proteins that appear dependent on EFP for proper expression. We show that these HGT events leave behind telltale genomic signatures and may have affected these species’ evolution, as many of these conserved polyproline motifs appear to be important for ATP binding and hydrolysis. These patterns of genome evolution suggest that horizontally transferred EFP and its modification systems interact with ‘native’ EFP with deleterious consequences.

## Results

### Genome dataset overview

Although EFP is an important protein in bacteria, as evidenced by its near universal conservation and the severe, detrimental effects of its loss^17,18,20,22,24^, it has been studied in only a handful of species^1,13–15,22,24^. The phylogenetic diversity of these species is limited mostly to the Gammaproteobacteria^1,14,22^. In order to investigate horizontal transfer of EFP from a wider diversity of species than purely experimental methods allow, we first needed to characterize these proteins. Presently, there are five known types of EFP. They differ in the amino acid residing at the tip of the conserved loop region in domain I, and the post-translational modifications of this amino acid. As detailed further in the Methods section, these comprise the lysine-EpmAB type EFP, the lysine-YmfI type, the unmodified lysine proline loop type, the arginine-EarP type, and the unmodified arginine-YeiP type^9^. We bioinformatically annotated all known EFPs within a dataset of more than 3000 bacterial species, comprising 35 phyla spread across 382 families. This procedure still left us with many ‘unknown’ EFP types—almost half of the genomes in our dataset (1606 genomes) encode an unknown type EFP.

To further characterize these unknown type EFP, we sorted all EFP sequences (3638 proteins) into ‘families’ using similarity network-based sequence homology clustering (*Methods*). In this process, we assigned protein sequences with ≥49% sequence identity and ≥80% sequence length alignment to the same family. We chose this threshold as a compromise between sorting known modification types into unique families and minimizing the number of families with few members due to poor representation of some clades in the dataset. This procedure left us with 15 EFP families (**Figure S1**). The largest of these families contained 2979 EFPs, including all lysine type EFPs—the EpmAB type (13.9%), the YmfI type (6%), the unmodified proline loop type (15.2%), and many EFPs of unknown modification (51%). The next largest families contained 299 EFPs primarily of the arginine-EarP modification type, and 265 EFPs of the arginine-YeiP type. The remaining families were unique to discrete phylogenetic clades. For example, we found the phyla Spirochaetota, Acidobacteriota, and Verrucomicrobiota to have particularly rich EFP diversity, encoding seven unique EFP types between them (**Figure S1 &** *Supplemental results*).

We found that the vast majority of bacterial genomes (2910 genomes, or 88.8% of our dataset) contain just one copy of the *efp* gene. A small proportion encoded two copies of the *efp* gene (362 genomes, or 11.0%), and just one species encoded three copies (the Gammaproteobacteria *Marinobacterium rhizophilum*). When two *efp* genes are present in a genome they usually encode different amino acids at the tip of the conserved loop region (314 or 86.5% of genomes with two EFPs encode EFPs with different amino acids at this position). Furthermore, the two EFPs are usually quite distinct in sequence composition (In 301 or 83% of genomes encoding two EFPs these EFPs fall into different sequence cluster families). In those species encoding two *efp* genes, the identity of the pair is usually a lysine-EpmAB type EFP and an arginine-YeiP type EFP (256 genomes, or 70.7% of genomes encoding two EFP). This pairing is most common among the Gammaproteobacteria (245 genomes), but also occurs in the phyla Desulfobacterota (5 genomes) and Planctomycetota (6 genomes). We found several other combinations of EFP pairs, which are detailed in the *Supplemental results*.

### Horizontal transfer of EFP

As a first pass at identifying horizontally transferred EFPs, we superimposed the bioinformatically inferred EFP modification types onto the phylogenetic tree of our bacterial species (**Figure 1**). Because EFP modification types evolved in distinct phylogenetic clusters^12,16^, this procedure can help identify EFP types that appear outside of the cluster where they originated. For example, the EarP rhamnose modification system originated within the Betaproteobacteria^16^. Any instances of the EarP type EFP residing outside this phylogenetic group therefore indicate HGT of *efp*. Using this approach, we identified 10 likely instances of EFP transfer, which are shown individually in **Figure 1**. They include 6 transfers of EpmAB type EFPs and 4 transfers of EarP type EFP. We see no evidence of transfer of the aminopentanol YmfI EFP type, perhaps because three separate genes are required for the synthesis and attachment of this modification^12^. We decided to investigate two instances of EFP transfer within the phyla Thermotogota and Planctomycetota further. We chose the Planctomycetota specifically because, as mentioned in the Introduction, they do not encode an otherwise universally conserved proline triplet in Valine tRNA synthetase^26^. Otherwise, these phyla had sufficiently many genomes (>20) to reliably identify HGT, and also comprise free-living bacteria unlikely to be undergoing genome degradation, a confounding phenomenon that could complicate our analyses.

**Figure 1:**
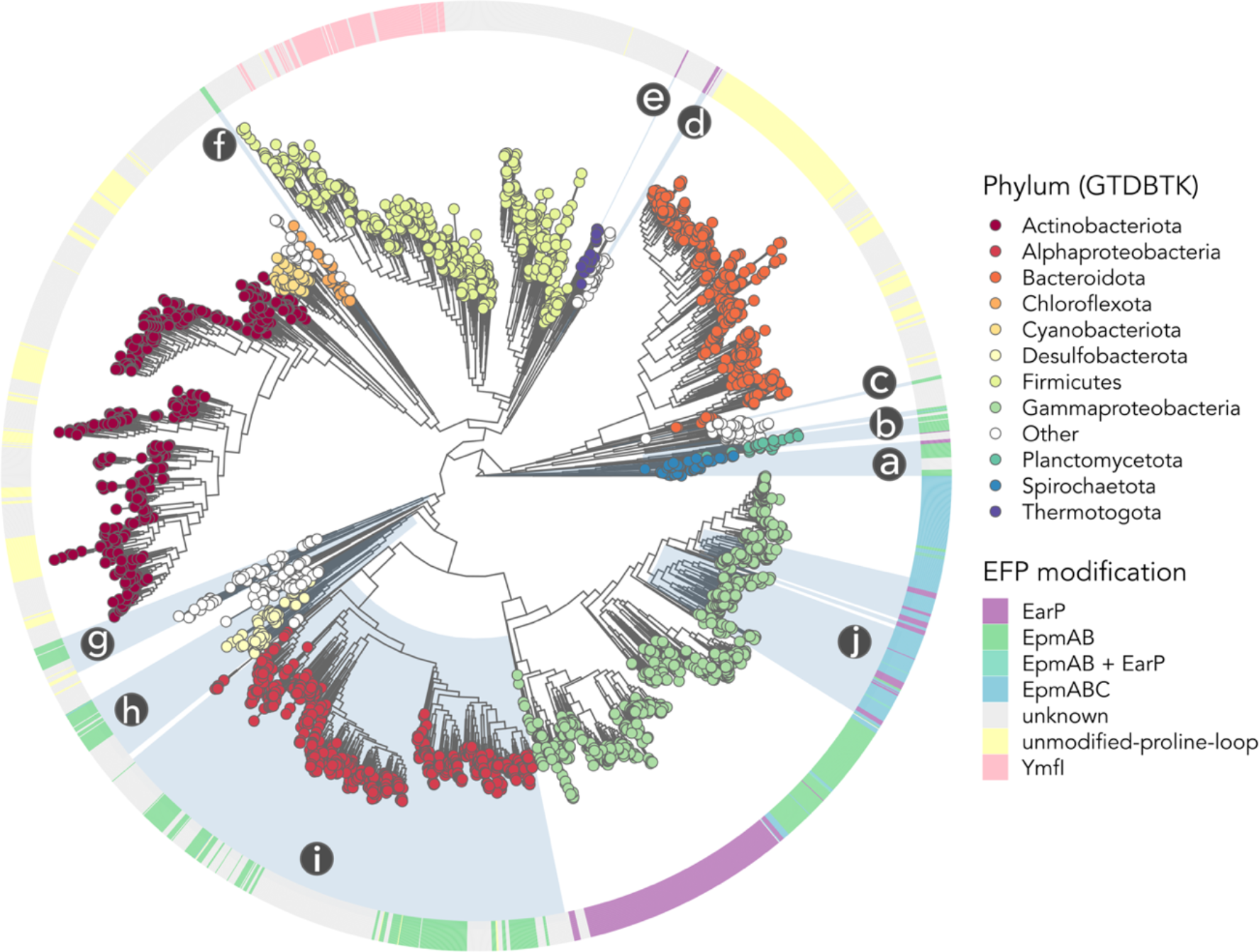
EFP modification systems evolved in phylogenetically conserved clusters of bacteria, making horizontal transfer of different *efp* types clearly discernable. We created this phylogenetic tree using the amino acid sequences of concatenated conserved proteins from >3000 bacterial genomes from 35 phyla. The ring indicates which EFP post-translational modification type is predicted to be encoded by each genome (*Methods*). The color of the circle at the leaves indicates which phylum each EFP sequence came from according to the Genome Taxonomy Database (GTDB)^29^. For clarity, only 11 phyla are colored, all others are termed ‘other’. (We note that Betaproteobacteriales is technically classified as an order within the Gammaproteobacteria in the GTDB^29^). Likely cases of EFP transfer are highlighted directly on the tree with blue shading and letters, and correspond to: a) Classes Leptospirae and Spirochaetia (EpmAB EFP) & family Treponemataceae (EarP EFP), b) Phyla Planctomycetota and Verrucomicrobiota (EpmAB EFP) & genus Planctopirus (EarP EFP), c) Family Fibrobacteraceae (EpmAB EFP), d) Family Fusobacteriaceae (EarP EFP), e) Family Petrotogaceae (EarP EFP), f) Family Dehalococcoidaceae (EpmAB EFP), g) Phyla Aquificota and Campylobacterota (EpmAB EFP), h) Members of the phyla Desulfobacterota, Desulfuromonadota, and Myxococcota (EpmAB EFP), i) Many members of the Alphaproteobacteria (EpmAB EFP), j) Gammaproteobacteria orders Pseudomonadales, Enterobacterales, Thiomicrospirales, and Thiotrichales (EarP EFP). In several genomes both EarP and EpmAB type EFPs coexist, i.e., in *L. lipolytica, Z. antarctica, M. rhizophilum*, and the genus *Plasticicumulans*).

### EFP transfer leads to loss of conserved polyproline motifs in the phylum Thermotogota

The phylum Thermotogota consists of mostly thermophilic, fermentative anaerobes with a distinctive “toga”-shaped outer cell envelope^27^. Members of the phylum are commonly found within hydrothermal vents, petroleum reservoirs, and hot springs^27^. Our phylogenetic tree indicates that an *efp* gene may have been horizontally transferred into this phylum (**Figure 1**). Specifically, while the majority of Thermotogota species encode an EFP type of unknown modification which falls within the most common EFP family (**Family 1, Figure S1**), two species from the Petrotogaceae family (*Geotoga petraeae* and *Oceanotoga teriensis*) encode the EarP type EFP and its modification system EarP (**Figure 1 & Figure S1, Family 3**). We used gene synteny (**Figure 2**) and gene-species tree phylogenetic distance comparisons (**Figure S2**) to verify the exogenous origin of this EarP type EFP. These analyses showed that the gene synteny of the EarP type EFP is inconsistent with other members of the phylum (the conserved placement of EFP within an operon together with the genes *pepP, yloU*, and *nusB* is lost, **Figure 2**) and that this EFP type is more sequence divergent from other Thermotogota EFPs than their shared phylogeny would predict (**Figure S2**). For example, the EarP type EFP of *O. teriensis* is more similar in sequence composition to EFPs from the Gammaproteobacteria family Burkholderiaceae (49.7% aa identity to *P. granuli* EFP) than to EFPs within its same phylogenetic family (39.5% aa identity to intrafamily member *M. hydrogenitolerans* EFP). Unexpectedly, these analyses uncovered a second case of EFP transfer within the same family. That is, species within the genera Petrotoga and Defluviitoga have also lost the conserved gene synteny present in all other Thermotogota (**Figure 2**), and as opposed to the preceding example, encode EFPs that are more similar in sequence to EFP from the Thermotogaceae family than their shared phylogeny would predict (**Figure S2**). For example, the EFP of *Defluviitoga tunisiensis* shares 75.1% aa identity to the EFP of *Thermotoga* sp. RQ7 of the Thermotogaceae family, yet this EFP shares only 61.1% aa identity to the EFP of intrafamily member *M. hydrogenitolerans*. It appears that species within the genera Petrotoga and Defluviitoga now encode an unknown type EFP that originated from a different family within Thermotogota, the Thermotogaceae.

**Figure 2:**
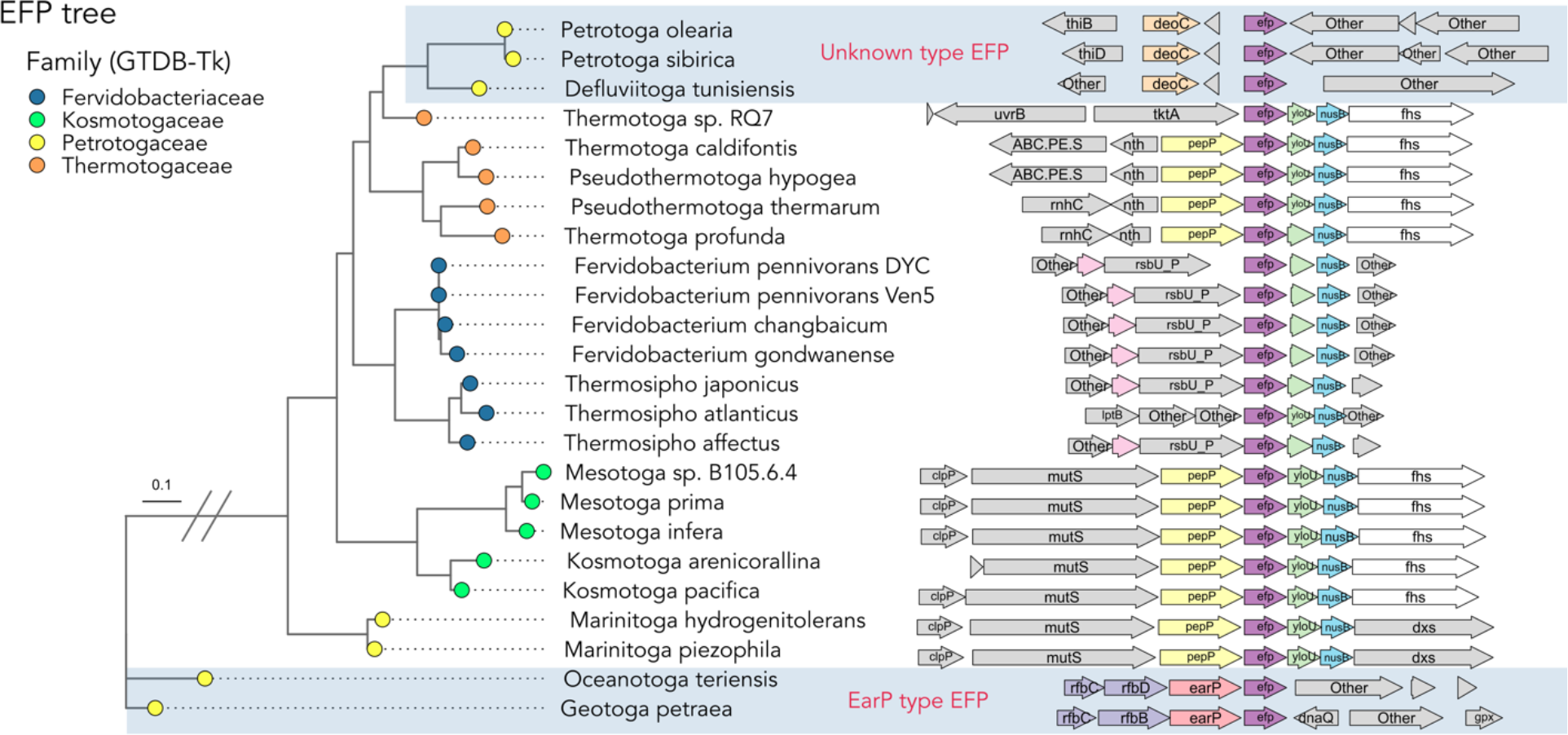
Horizontal gene transfer of EFP in the phylum Thermotogota. **Left:** Phylogenetic tree of EFP amino acid sequences in the phylum Thermotogota (*Methods*). The phylogeny of EFP in the family Petrotogaceae is not congruent with the overall phylogenetic tree of the family’s species (**Figure S2**). Within the Petrotogaceae family two separate gene transfer events of the *efp* gene have occurred (highlighted with blue grey shading across the figure), one from within the Thermotogaceae family (top) and one from outside the phylum (bottom). The color of the circle at the tree’s tips represents the family these genomes belong to, according to the Genome Taxonomy Database (GTDB)^29^. The tree branch with the indicated scale break has been shortened to 1/3 of its original length. **Right:** Gene synteny of each EFP gene from the tree on the left. For clarity, only select genes of interest are indicated with color. The ‘native’ copy of EFP (purple, centered) within the Thermotogota consistently co-occurs with YloU (mint green) and NusB (blue). This high conservation of synteny is absent for the EFPs highlighted within the blue-grey boxes, which supports their horizontal transfer into the corresponding genomes. Additionally, the EarP type EFPs have been transferred together with their modifying enzyme (EarP, salmon), and with several genes in the rhamnose biosynthesis pathway (light purple).

To investigate the effect of horizontal transfer of EFP on polyproline motifs in the Thermotogota, we first clustered all proteins within the phylum into families based on amino acid sequence identity (*Methods*), then identified conserved polyproline motifs and polyproline motif-containing proteins within these families. We next performed phylogenetic ANOVAs^28^ to test the null hypothesis that horizontal transfer of EFP is not associated with a loss of conserved polyproline motifs, or of the proteins which contain them, while taking the phylogeny of our species into consideration. We found that horizontal transfer of EFP is significantly associated with the loss of 11 polyproline motifs or polyproline motif-containing proteins (**Figure 3**). Loss of entire polyproline motif-containing proteins may indicate that the polyproline motif is crucial to their function, as is the case for the polyproline motif in the glucose importer component EII^Glc^ of *Corynebacterium glutamicum*^25^.

**Figure 3:**
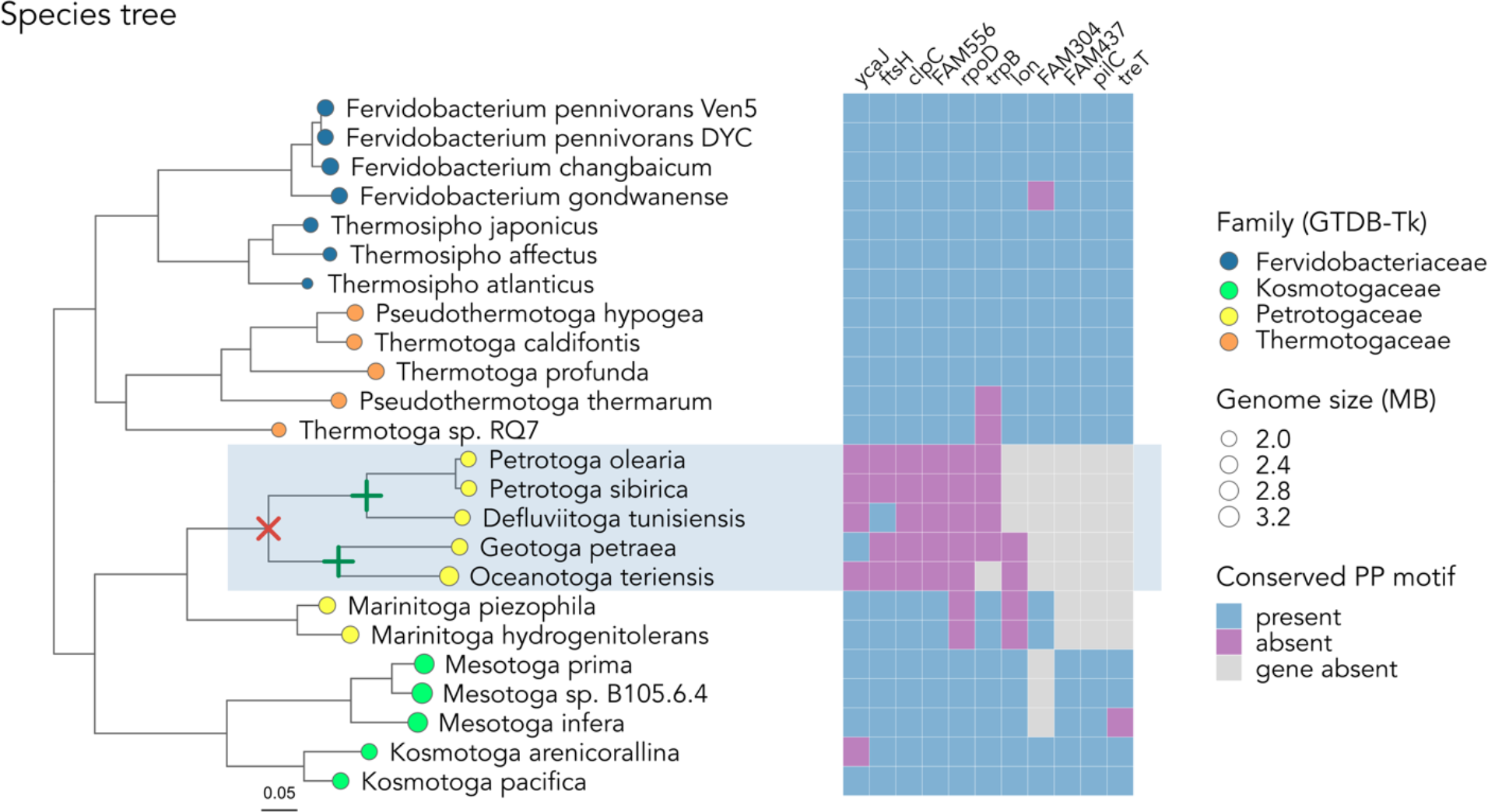
Eleven polyproline motifs and polyproline motif-containing proteins that are significantly associated with horizontal transfer of EFP into the Petrotogaceae family. We built this phylogenetic tree using amino acid sequences of 43 concatenated and conserved marker genes generated by CheckM^28^ (details in *Methods*). The color of the circle at the tree’s tips represents the family these genomes belong to, according to the Genome Taxonomy Database (GTDB)^29^. The size of the circle corresponds to relative genome size. Loss of ‘native’ EFP is indicated with a red X on the phylogenetic tree, while gain of a horizontally transferred EFP is indicated with a green +. The exact timing and order of these events is unknown. Species encoding horizontally transferred EFP are highlighted with grey-blue shading across the figure. The heatmap on the right shows for eleven proteins the presence of a conserved polyproline motif (blue), the absence of the polyproline motif in the same position of the protein (purple), or the complete absence of the protein (grey). The eleven proteins are, from the left-most to the right-most column, as indicated by their acronyms: The putative ATPase YcaJ, the cell division protease FtsH, the ATP-dependent Clp protease ATP-binding subunit ClpC, the cation efflux protein FAM556, the polymerase primary sigma factor RpoD, the tryptophan synthase beta chain TrpB, the ATP-dependent Lon protease, the uncharacterized B12-binding /radical SAM-type protein FAM304, the uncharacterized metalloprotease FAM437, the type IV pilus assembly protein PilC, and the trehalose synthase TreT. Proteins with the prefix FAM-are not annotated by the KEGG database, and this designation refers to their Silix^37^ identifier (*Methods*).

Next, we examined the polyproline motif-containing proteins that were significantly associated with horizontal transfer of EFP within members of the Petrotogaceae. These proteins were functionally diverse. They include three ATP-dependent proteases (Lon, ClpC, FtsH), two enzymes related to metabolism (the tryptophan synthase beta chain TrpB, the trehalose synthase TreT), four poorly characterized proteins (the putative ATPase YcaJ, a cation efflux protein annotated as FAM556 by our amino acid sequence clustering (*Methods*), an uncharacterized B12-binding /radical SAM-type protein FAM304, an uncharacterized metalloprotease FAM437), and two proteins of varied functions (type IV pilus assembly protein PilC, RNA polymerase primary sigma factor RpoD). Many of the polyproline motifs these proteins contain are highly conserved among our 3000 bacterial genomes (**Table S1**). For example, the polyproline motifs located within the bacterial proteases are generally highly conserved among bacteria—85.5%, 97.9%, and 98.8% of FtsH, ClpC, and Lon proteins contain polyproline motifs at these exact positions, respectively. This strong conservation implies that these polyproline motifs are important to their resident proteins and that these proteins likely depend on EFP for expression.

### EFP transfer leads to loss of conserved polyproline motifs in the phylum Planctomycetota

Next, we examined the suspected horizontal transfer of EFP and its effects on members of the phylum Planctomycetota. In the course of a previous study^3^, we discovered that the loss of a polyproline triplet in the protein Val tRNA synthetase, which is otherwise conserved across all three domains of life^26^, coincided with a case of horizontally transferred EFP in this phylum. We wanted to find out if additional patterns of polyproline motif loss coincided with this event. The phylum Planctomycetota includes cosmopolitan species that can be found in soil, aquatic, and wastewater habitats^29^. These bacteria possess diverse cell structures and metabolism. Some species divide by budding, some have cytoplasmic compartments, some perform anaerobic ammonium oxidation, and many have complex life cycles featuring transitions between sessile and swimming states^29^. Our initial analyses (**Figure 1**) indicated that Planctomycetota species have undergone at least three EFP transfer events. As a result, they now encode at least four distinct types of EFP, including EFP types with unknown modifications (likely the ‘native’ EFP types, which cluster into three distinct EFP families, **Figure S1**), EpmAB type EFP, EarP type EFP, and YeiP type EFP.

As before, we verified HGT events using gene synteny (**Figure S3**) and phylogenetic distance comparisons between gene and species trees (**Figure S4**). We found that members of the class Planctomycetes, including the families Planctomycetaceae, Pirellulaceae, and Thermoguttaceae, no longer encode the native EFP type present at the base of the Planctomycetes tree, and instead encode an EpmAB type EFP (**Figure S3**). This EFP type is distinct in sequence to all other types in the phylum (**Figure S4**), and in most species is flanked by the EpmB gene (**Figure S3**). A different, EarP type EFP, is encoded by members of the genus Planctopirus, and is flanked by the EarP gene (**Figure S3**). The final horizontally transferred EFP type, YeiP is encoded somewhat sporadically within the class Planctomycetes, including within the Pirellulaceae, Isosphaeraceae, and Thermoguttaceae families (**Figure S3**). Consistent with its identity as a “secondary” EFP subfamily^14^, the YeiP type EFP is in all species paired with a non-YeiP type EFP.

Following the same procedure as for the phylum Thermotogota, we found that three polyproline motifs were significantly associated with horizontal transfer of EFP, but in two different subsets of the class Planctomycetes (**Figure 4**). We found that all members of the class Planctomycetes that no longer encoded the native Planctomycetota EFP also lost the polyproline motif containing Lon protease, similar to what we had observed in the Thermotogota (**Figure 3**). Additionally, all members of the class Planctomycetes lost the polyproline-motif containing protein isoleucine tRNA synthetase type 1, instead encoding isoleucine tRNA synthetase type 2. Two distinct forms of isoleucine-tRNA synthase exist among bacteria—the typical form present in *E. coli* and most other species (type 1) and a second form more closely related to eukaryotic type IleS, which lacks tRNA-dependent pre-transfer editing activity (type 2)^41^. In just the family Planctomycetaceae, we found that the loss of conserved polyproline motifs in valine tRNA synthetase was associated with EFP transfer (**Figure 4**).

**Figure 4:**
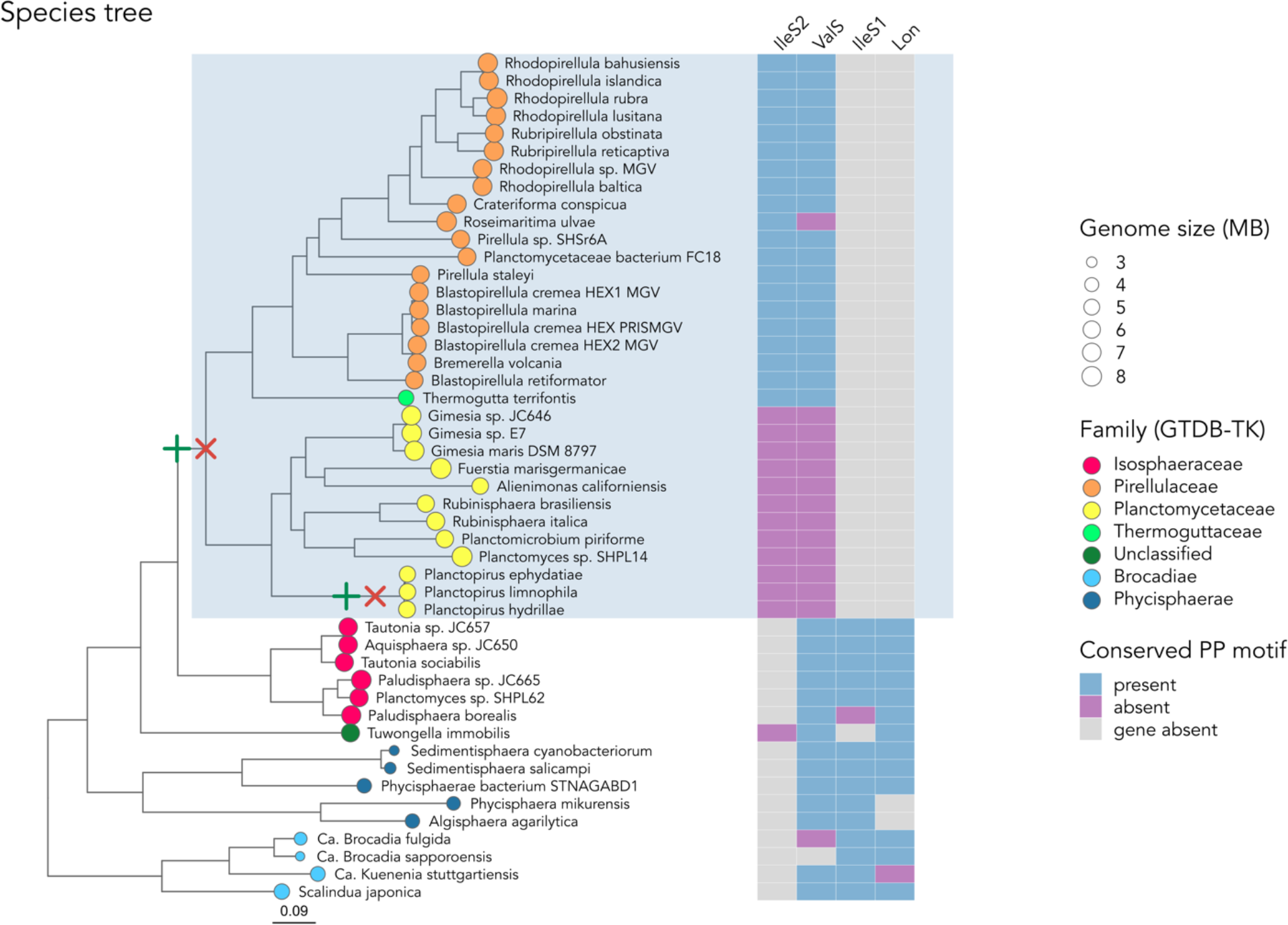
Three polyproline motifs and polyproline motif-containing proteins are significantly associated with horizontal transfer of EFP in the class Planctomycetes. We built this phylogenetic tree using amino acid sequences of 43 concatenated and conserved marker genes generated by CheckM^28^ (details in *Methods*). The color of the circle at the tree’s tips represents the family these genomes belong to, according to the Genome Taxonomy Database (GTDB)^29^. The size of the circle corresponds to relative genome size. Loss of native EFP is indicated with a red X on the phylogenetic tree, while gain of horizontally transferred EFP is indicated with a green +. The exact timing and order of these events is unknown. Species encoding horizontally transferred EFP are highlighted with grey-blue shading across the figure. The heatmap on the right reports the presence of a conserved polyproline motif (blue), the absence of the polyproline motif in the same position in the protein (purple), or the complete absence of the corresponding protein (grey). Sequence homology clustering of Planctomycetes proteins revealed two distinct forms of isoleucine-tRNA synthase, the typical form present in *E. coli* and most other bacteria (type 1) and a second form more closely related to eukaryotic type IleS which lacks tRNA-dependent pre-transfer editing activity (type 2)^41^. While IleS2 was not significantly linked with horizontal transfer of EFP after phylogenetic correction, its appearance in the phylum Planctomycetota and loss of a highly conserved polyproline motif led us to include it here. The four proteins displayed are, in order: isoleucine tRNA synthetase type 2 IleS2, valine tRNA synthetase ValS, isoleucine tRNA synthetase type 1 IleS1, and ATP-dependent Lon protease.

### Characteristics of proteins sensitive to horizontal transfer of EFP

From our investigations of the phyla Thermotogota and Planctomycetota, two major groups of proteins that appear to be sensitive to EFP transfer emerged. These are ATP dependent proteases (Lon, ClpC, and FtsH) and class I tRNA synthetases (ValS, IleS1, and IleS2). We searched for commonalities within these groups of proteins and tried to determine whether the polyproline motifs linked to EFP transfer are important for the function of these proteins.

The polyproline motifs within the three proteases (Lon, ClpC and FtsH) are well conserved among our wider dataset of over 3000 bacterial species. Specifically, 98.8% of Lon, 97.9% of ClpC, and 85.5% of FtsH proteins have polyproline motifs in this position (**Table S1**). Upon further examination, we discovered that each of these well-conserved polyproline motifs are located within the same PFAM domain (PF00004). This domain characterizes a diverse ATPase family associated with a broad range of cellular activities^31^. Within this domain, each polyproline motif is positioned within the ATP binding pocket (**Figure 5**). Although not predicted to be a protease, we also found the same pattern in the putative ATPase YcaJ (**Figure 3**). Mutations of residues within this ATP binding pocket are known to inactivate Lon protease^32^ and FtsH^33^ in *E. coli*. Furthermore, the expression of Lon and Clp proteins is dependent on EFP in *E. coli* ^6,34^ and *Salmonella enterica*^1^. Expression of FtsH (known by the synonym HflB) is EFP-dependent in *Salmonella enterica*^1^. Together, these observations suggest that these highly conserved polyproline motifs are important for ATP binding and hydrolysis in ATP-dependent proteases. Consequently, these proteins are prone to depend on EFP for proper expression.

**Figure 5:**
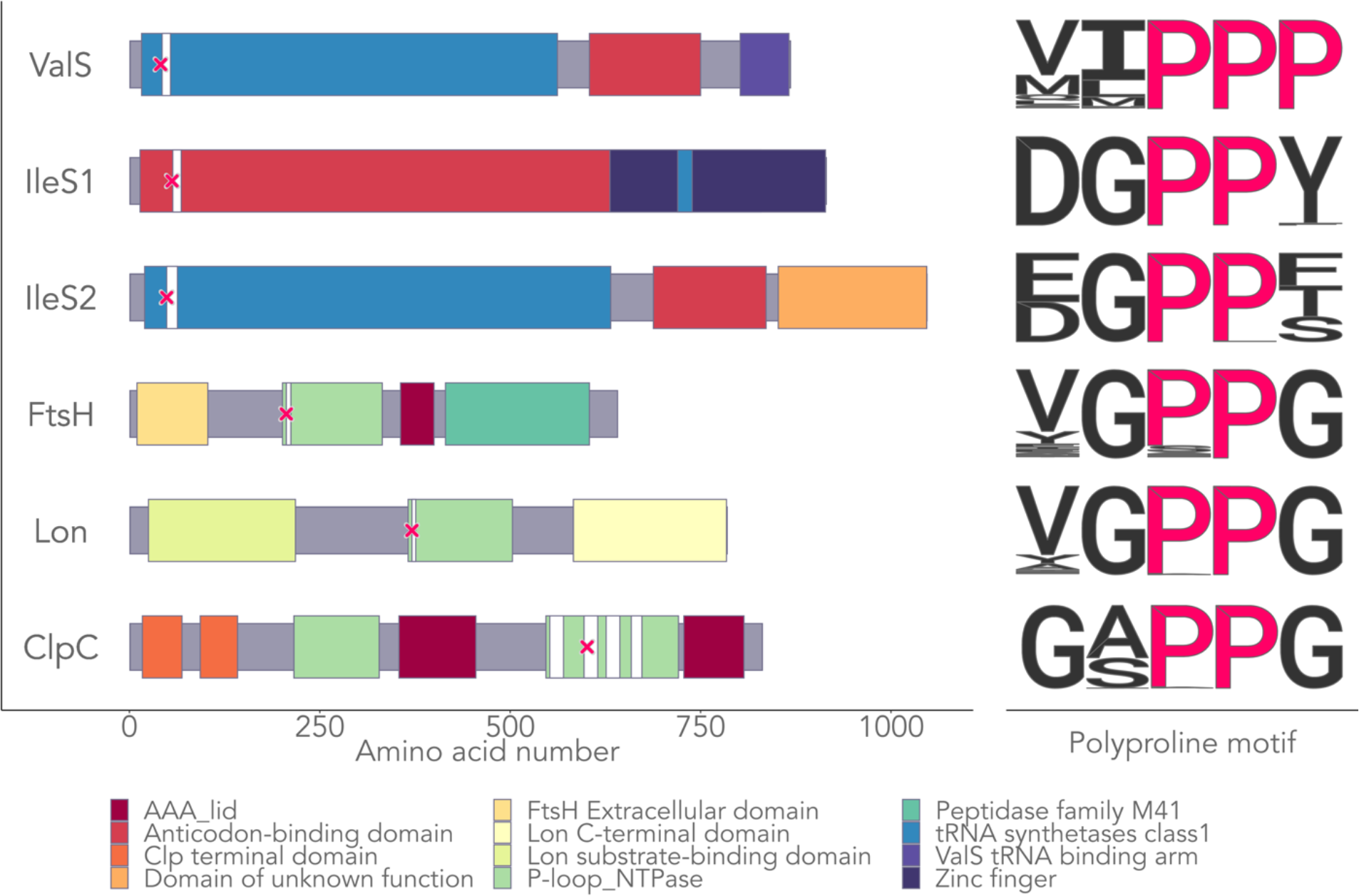
Polyproline protein sequence motifs sensitive to the loss of native EFP. **Left:** We found conserved polyproline (PP) motifs within three tRNA synthetases (ValS, IleS1, and IleS2) and three proteases (FtsH, Lon, ClpC) to be significantly linked to the loss of native EFP. That is, in species that no longer encode their phylogenetically ‘native’ EFP, these motifs are often absent. Protein length is indicated by grey narrow horizontal bars, on top of which PFAM domains annotated by InterProScan^50^ are displayed, colored by either their PFAM or PFAM clan description^34^. Locations of conserved PP motifs are indicated with a pink X. White vertical bars correspond to regions predicted to interact with ATP. More specifically, they represent (i) the conserved tRNA class I His-Ile-Gly-His “HIGH” consensus motifs^46^ InterproID (IPR001412) for the tRNA synthetases, and (ii) ATP binding regions (InterproID IPR001270 or PIRsitepredict ID PIRSR001174-2) for the proteases. For consistency, all proteins shown in the figure are from the genome of *Kosmotoga arenicorallina* (which encodes all indicated PP motifs and the native EFP of the phylum Thermotogota), except for IleS2, which is from *Mesotoga prima*. (*K. arenicorallina* encodes only IleS1, not IleS2. *M. prima* also encodes the native EFP type of the phylum Thermotogota). **Right:** Sequence logos of the well conserved PP motifs represented with a pink X in panel A) within our dataset of >3000 bacterial genomes from 35 phyla. Specifically, 99.8% of ValS, 99.7% of IleS1, 99.3% of IleS2, 85.5% of FtsH, 98.8% of Lon, and 97.9% of ClpC proteins in our dataset have PP motifs in the indicated position. The consensus motifs of the species which do not encode the canonical PP motifs are, following the figure order from top to bottom: “MIPLP”, “DGPIY”, “DGPIT”, “VGSPG”, “VGAPG”, and “GSAPG”.

The polyproline motifs within the three class I tRNA synthetases (ValS, IleS1, and IleS2) are the most strongly conserved in our wider 3000 species dataset. Specifically, 99.8% of ValS, 99.7% of IleS1, and 99.3% of IleS2 proteins have polyproline motifs in this position. Indeed, a previous study found that these polyproline motifs are invariant across all domains of life^26^. The ValS polyproline motif is encoded by all non-Planctomycetaceae species in our wider genome dataset, with only one exception, the Cyanobacterium *Pseudanabaena sp*. PCC 7367. However, all other available genomes from the Pseudanabaenaceae family encode the polyproline motif, meaning there are no additional genomes that can help validate the accuracy of this exception. Similar to the ATP-dependent proteases, the polyproline motif in ValS lies within the ATP binding region of this protein, near the His-Ile-Gly-His (“HIGH”) motif that is characteristic of this class of tRNA synthetases (**Figure 5**)^26,35^. Mutation of the polyproline motif in *E. coli* results in a ValS that nonproductively hydrolyzes ATP to ADP and mischarges tRNA^Val^ with threonine^26^. The polyproline motifs of IleS type 1 and 2 also lie next to the HIGH motif of these proteins (**Figure 5**). While the function of this motif in IleS has not been characterized, it is also likely to be involved in ATP binding and hydrolyzation.

Expression of ValS is strongly dependent on EFP in *E. coli*^6,26^ and *S. enterica*^1^. Indeed, under-expression of ValS in *E. coli* EFP deletion mutants is partially responsible for their growth defect^26^.

In summary, we find that the highly conserved polyproline motifs present in both the ATP dependent proteases and class I tRNA synthetases are likely involved in ATP binding. Many of these proteins are dependent on EFP for normal expression across multiple species^6,26^. The alteration of these highly conserved polyproline motifs, or the loss of the proteins containing them, coincides with EFP transfer in two independent phyla. We conclude from these observations that EFP transfer is associated with a disruption of global EFP function. In response to this disruption, selective pressure against polyproline motifs emerges, namely in proteins that are dependent on EFP for proper expression.

## Discussion

Loss of EFP activity can have dramatic impacts upon bacteria, leading to the under-expression of proteins that contain polyproline motifs^17,18,20,22^, general growth defects ^8,17–20^, and in some species cell death^22,24^. During the course of a previous analysis^3^, we discovered hints of a connection between the horizontal transfer of *efp* and disruption of EFP activity through the loss of highly conserved polyproline motifs. In this work, we thoroughly investigate this connection, and study how EFP transfer may have affected modern-day bacterial species from a wider phylogenetic diversity than experimental methods allow. To this end, we first characterized EFP from a set of over 3000 bacterial genomes from 35 phyla. We found that horizontal transfer of the *efp* gene has occurred multiple times along the bacterial tree of life, with several cases of EarP type EFP and EpmAB type EFP found outside their clades of phylogenetic origin (**Figure 1**). We examined in more detail two phyla whose members contain horizontally transferred EFP, the Thermotogota (**Figure 2 & S2**) and the Planctomycetota (**Figure S3 & S4**) and found that EFP transfer is consistently associated with the loss of polyproline motifs and polyproline motif-containing proteins (**Figure 3 & 4**). In particular, we found two groups of proteins which seem to be particularly sensitive to HGT of *efp*. These are the ATP dependent proteases Lon, ClpC, and FtsH, as well as the class I tRNA synthetases IleS (type 1 and 2) and ValS (**Figure 5**). We show that the position of the polyproline motifs within these proteins is highly conserved within the wider diversity of our 3000 bacterial species (**Table S1**). That is, 85.5-99.8% of these six proteins contain polyproline motifs in the same conserved positions. Additionally, these motifs are all within or in close proximity to ATP-binding domains (**Figure 5**). While the polyproline motif within ValS is known to be crucial for proper ATP hydrolyzation^26^, the precise function of the other polyproline motifs have not been studied in detail.

Our observations suggest that genomes which encode horizontally transferred *efp* at one point experienced a disruption in global EFP activity. This disruption would have negatively impacted the expression of polyproline motif containing proteins, as has been shown experimentally in multiple species^17,18,20,22^. To mitigate this disruption, some proteins seem to have lost highly conserved polyproline motifs, thereby severing their dependence on EFP for expression. In other proteins, like Lon protease, these highly conserved polyproline motifs may be critical to the function of the protein, leading instead to the loss of the entire protein. However, with only present-day genomes to work from, we can merely hypothesize what the cause of this disruption was. One possibility is that species within the Planctomycetota and Thermotogota first lost their ‘native’ form of EFP, which would make EFP transfer especially advantageous. While the present-day Thermotogota and Planctomycetota species which encode horizontally transferred versions of EFP no longer encode the EFP native to their phylum (**Figure 2 & S3**), it is relevant to note that we do find cases of horizontally transferred EFP coexisting with native EFP, albeit in only a small fraction of our overall dataset (8 genomes from the Desulfuromonadota, 3 genomes from the Chloroflexota, and 25 genomes from the Alphaproteobacteria). Additionally, EFP loss has not been observed in free-living organisms: EFP is universally conserved in bacteria, with the exception of a few endosymbiotic species undergoing genome degradation^9,36^. EFP loss can have stark consequences, as experimental EFP loss is known to be lethal in two species^22,24^, and cause severe phenotypes in other tested species^8,17–20^.

Another possibility is that the transfer of an extraneous EFP and its modification system into the Thermotogota and Planctomycetota happened before any loss of native EFP. In this case, deleterious interactions between the native and horizontally transferred proteins may have perturbed the expression of EFP-dependent proteins. It is tempting to speculate that these perturbations arose from deleterious interactions between the horizontally transferred modification system and the native EFP. Different EFP types and their corresponding modification systems co-evolved, such that switching just the conserved, post-translationally modified amino acid of an EFP is not sufficient to switch its modification system^14,16^. For example, the function of an EarP-type EFP_R32K_ from *Shewanella oneidensis* cannot be rescued by expression of the EpmAB system from *E. coli*, and neither can the function of an EpmAB-type EFP_K34R_ from *E. coli* be rescued by the expression of EarP from *S. oneidensis*^14^. Indeed, subjecting an EFP to a non-native post-translational modification can inhibit its function below that of the post-translationally unmodified state^16^. If a non-native modification system can modify and impair the function of the native EFP type, overall EFP function could be compromised, leading to the genome evolution patterns we observe. In support of this hypothesis, we note that the conserved positions within EFP that bear post-translational modifications in both the Thermotogota and the Planctomycetes are theoretically compatible with their horizontally transferred modification systems. That is to say, the native EFP within the Thermotogota is an arginine type EFP, and the horizontally transferred EarP modifies arginine residues (**Figure S5**). Likewise, the native EFP within the Planctomycetota fluctuates between a lysine and asparagine type (**Family 11, Figure S1**), and the horizontally transferred EpmAB system modifies lysine residues (**Figure S6**).

Lastly, we note that our two focal phyla in which the horizontal transfer of *efp* occurred have very different lifestyles. The Petrotogaceae are fast-growing thermophiles with optimal growth temperatures between 45 - 60°C^27^. The Planctomycetota are mostly mesophilic and notoriously slow growing, with doubling times ranging from 6 hours to 1 month^37^. In a previous study, we found that both slow-growing bacteria and thermophilic bacteria are enriched in polyproline motifs when compared to fast-growing mesophiles^3^. We hypothesized that this is because of lower selective pressures on translation speed in slow-growing organisms, which do not need to synthesize proteins rapidly, and the catalytic “boost” that thermophiles derive from high growth temperatures, which can lead to naturally higher rates of translation^3^. If so, these two groups of organisms may be particularly well poised to endure disruptions in EFP function, which may give them sufficient time to adapt evolutionarily by altering polyproline motifs to make key proteins less EFP-dependent. Tentative support for this hypothesis comes from studies of the *efp* deletion mutant phenotype in *E. coli* ^8^. The growth defect of these mutants is less severe when *E. coli* cells grow more slowly, indicating that dependence on EFP is strongest when protein expression and demand for translational efficiency is high^8^. However, these speculations require experimental validation, as all EFP mutants thus far have been studied in fast-growing, mesophilic organisms^8,17–20^.

The power of comparative genomics comes from the ability to leverage the evolutionary history of thousands of species, and to make predictions based upon their signatures of genomic change. It has allowed us to discover a recurrent connection between the horizontal transfer of EFP and signs of EFP dysfunction through the loss of highly conserved polyproline motifs and polyproline-motif containing proteins. Ancient disruptions in EFP activity have not only left clear traces in the genomes of modern-day species, but they may also have impacted the evolution of these species to the present day.

## Materials & Methods

### Selection of Study Phyla

We used a set of 3265 phylogenetically diverse bacterial genomes that we characterized in a previous study^3^. These genomes span 35 phyla and were selected to maximize phylogenetic diversity—we included only one genome per Average Nucleotide Identity cluster, or in other words, only one genome per species present in the Integrated Microbial Genomes (IMG) database^39^. We checked each genome for completeness and contamination with CheckM^40^ (with cutoffs of ≥90% completeness and ≤5% contamination), and reassigned taxonomy using the Genome Taxonomy Database and GTDB-Tool kit (GTDB-Tk) version 0.2.2^41^.

We used these genomes to identify taxonomic groups that may have lost their ‘native’ EFP and obtained exogenous EFP through horizontal gene transfer, as identified through ‘non-native’ types of post-translational EFP modifications. The presence of such modifications can be computationally inferred by identifying the genes that encode the proteins performing the modification, as well as by identifying the modified amino acid within EFP. Because EFP modification types originated in distinct phylogenetic clusters of bacteria, EFP modification types that occur outside their cluster of origin indicate that an EFP-coding gene and its associated modification genes have been transferred.

To detect such instances of HGT we first created a phylogenetic tree of all our genomes using 43 concatenated and conserved marker protein sequences generated by CheckM (v1.0.12)^40^ for a previous study^3^, then used IQ-TREE (v1.6.12)^42^ to build the tree. We used the model finder feature^43^ included in IQ-TREE to determine the best-fit substitution model for our tree (which was the “LG+F+R10” model). This tree is shown in **Figure 1** and is rooted with the genome of the Archaeon *Haloquadratum walsbyi*.

Next, we determined the post-translationally modified amino acid residue of each EFP. We did this by first aligning all proteins annotated as EFP by the KEGG functional database (Kegg Orthology term: K02356) using MUSCLE (v3.8.31)^44^ with default settings. We then identified the post-translationally modified amino acid residue using the MUSCLE alignment with validated EFP sequences as a guide. We assigned EFP modification types by searching for a mix of annotations from the Clusters of Orthologous Groups (COG)^45^ and Pfam (Protein families) databases^46^. First, we considered a species to have an EFP modified by β-(*R*)-lysylation if its genome encoded the genes for EpmA (COG2269) and EpmB (COG1509). To identify the optional hydroxylation of EpmAB-modified EFP, we looked for the gene encoding EpmC (pfam04315)^11^. Second, we considered an EFP to be modified by rhamnosylation if its genome encoded EarP (pfam10093)^14^. Third, we considered an EFP to be modified by the 5-aminopentanol moiety if its genome yielded BLASTP (v2.13.0+)^47^ hits in a search for the three proteins that carry out the attachment (YmfI, YnbB, and GsaB)^13^, using *Bacillus subtilis* orthologs of these proteins as our query sequences. We used different bit-score cut-offs for each gene (YmfI : 145, YnbB : 200, and GsaB : 525). We used a BLAST-based approach for these proteins because they do not have a consistent annotation in either the COG or Pfam databases. Fourth, we identified YeiP type EFPs using the TIGRfam (The Institute for Genomic Research Protein Families) database^48^. TIGRfam is a collection of manually curated protein families similar to Pfam, and it is the only database that distinguishes between canonical EFP (TIGR00038) and the YeiP (TIGR02178) subtypes. We validated these four EFP modification type assignments by confirming that the respective *efp* genes encoded lysine (EpmAB and YmfI/YnbB/GsaB) or arginine (EarP and YeiP) at the conserved modification position. Lastly, we identified EFPs that are thought to function without any modification by searching for the characteristic unmodified proline loop in the conserved positions 30 and 34 (P_30_NNNP_34_) within EFP protein sequences^15^.

We used these assigned EFP types to identify which bacterial phyla to target further for in-depth investigation. Our initial findings led us to focus on the Planctomycetota, and we also chose the Thermotogota because they are a free-living clade unlikely to be undergoing genome degradation, and whose genomes are well-represented in our data set (with >20 genome sequences per phylum).

### Verifying HGT of the *efp* gene

Within these target phyla, we next constructed phylogenetic trees of the EFP protein sequence, as well as species trees to verify the exogenous origin of putative horizontally transferred EFPs. If an EFP-coding gene has been transferred horizontally, the species tree and the EFP tree will be discordant, because the transferred EFP gene has not evolved within the clade it now resides in. We built phylum specific EFP gene trees with IQ-TREE (v1.6.12) after aligning the EFP protein sequences using MUSCLE^31^. To prepare the species tree we used 43 concatenated and conserved marker protein sequences generated by CheckM (v1.0.12)^40^ for a previous study^3^, then used IQ-TREE to build the tree.

We verified the HGT of EFP with two methods. First, we compared the species and EFP trees by plotting the distance (cumulative branch length) between pairs of species on each tree. We found that generally, these cumulative branch lengths are highly correlated between the two trees. In other words, the rate at which amino acid substitutions occur in the native *efp* gene is similar to the rate of amino acid substitutions within the 43 conserved marker protein sequences we used to build the species tree. This association renders likely *efp* horizontal transfer events visually obvious outliers (**Figures S2 & S4**). Second, we investigated gene synteny between the *efp* gene and adjacent genes for each genome from our phyla of interest to verify that gene order differs between putatively transferred and native *efp* copies (**Figures 2 & S3**).

### Loss of conserved polyproline motifs

To find polyproline motifs whose loss coincided with horizontal transfer of EFP, we used a computational method that is independent of annotation databases. This is important, because our phyla of interest are poorly studied, and thus poorly annotated. (For example, 45% of genes in the Planctomycete *Planctopirus limnophila* have no predicted function). For each phylum we were interested in, we combined all proteins from each genome within that phylum into one file (49 species for the Planctomycetes and 24 species for the Thermotogota), then performed an all-versus-all BLASTP (v2.13.0+)^47^ search, using default parameters. With the resulting BLASTP output, we performed de novo clustering of the proteins using Silix (v1.3.0)^49^. Silix is a software tool that clusters protein sequences into homologous families using similarity networks^49^. Sequences with ≥35% sequence identity and ≥80% sequence alignment were assigned to the same family. This generated 2379 and 74409 protein families respectively, for the phyla Thermotogota and Planctomycetes. We aligned each family of proteins with MUSCLE and used custom python scripts to locate polyproline motifs within the proteins.

We tested the null hypothesis that horizontal transfer of the *efp* gene into a genome is not associated with 1) the loss of conserved polyproline motifs within particular proteins, or 2) the loss of entire proteins containing well conserved polyproline motifs. We sub-selected the proteins we tested using the following criteria. First, we created two groups of genomes—those which encoded a putative horizontally transferred *efp* gene (HGT group) and those which still encoded their native *efp* gene (non-HGT group). To select which conserved polyproline motifs within particular proteins to test, each polyproline motif and the protein it resides within must pass one of two conditions:

1. The protein and polyproline motif must be well conserved among the non-HGT group and poorly conserved among the HGT group (70% of genomes in the non-HGT group encode the protein and the polyproline motif is present at a conserved position in at least 85% of these, while genomes in the HGT group encode the polyproline motif less than or equal to 20% of the time).
2. The protein and polyproline motif must be well conserved among the non-HGT group, but the protein itself is rare in the HGT group (same cutoffs as above for the non-HGT group, but the polyproline containing protein is present in 20% or fewer genomes in the HGT group).

For polyproline motifs or polyproline containing proteins that passed these filters, we used the phylANOVA function from the R package phytools (v. 0.7-70)^28^ to test our null hypothesis that horizontal transfer is not associated with a loss of polyproline motifs or of proteins harboring them. PhylANOVA allowed us to control for the phylogenetic interdependence of protein sequences using the species trees described above. We corrected P-values for multiple testing using the Benjamini & Hochberg method at a false discovery rate of 0.1.

### Characterizing conservation of protein sites with polyproline motifs

For each annotated protein and/or polyproline motif whose loss was significantly associated with the HGT of *efp*, we also determined how conserved the relevant polyproline motif is in our wider 3000 species genome set. To this end, we first extracted the amino acid sequences of these proteins (YcaJ, FtsH, ClpC, RpoD, TrpB, Lon, PilC, TreT, ValS, and IleS) from each genome in our dataset using KEGG annotations. For each set of proteins, we again used Silix to cluster its homologues into families using the same parameters as described above. We clustered these protein sequences because we found some orthologous proteins to be annotated with the same KEGG Orthology term. For example, the Lon protease, the Lon-like protease BrxL, and the sporulation protease LonC are all annotated as ATP-dependent Lon proteases (K01338) by KEGG. We next aligned the Silix families bearing our protein of interest using MUSCLE, and then used our custom python scripts to locate conserved polyproline motifs. We then calculated how conserved each of these polyproline motifs are within each target protein across our 3000 genomes dataset (**Table S1**).

## Data availability

All genomes used in this study are publicly available from JGI’s IMG database^39^. R scripts and all files needed to reproduce these analyses and figures are available at: https://github.com/tessbrewer/pattern_project.

## Acknowledgements

We acknowledge funding from the European Research Council under Grant Agreement No. 739874, as well as from Swiss National Science Foundation grant 31003A_172887, and from the University Priority Research Program in Evolutionary Biology at the University of Zurich.

