## Supplemental materials for "Horizontal gene transfer of a key translation protein has shaped the polyproline proteome"

**Contains:**

Supplemental Figures 1-6, Supplemental Results, and Supplemental Table 1

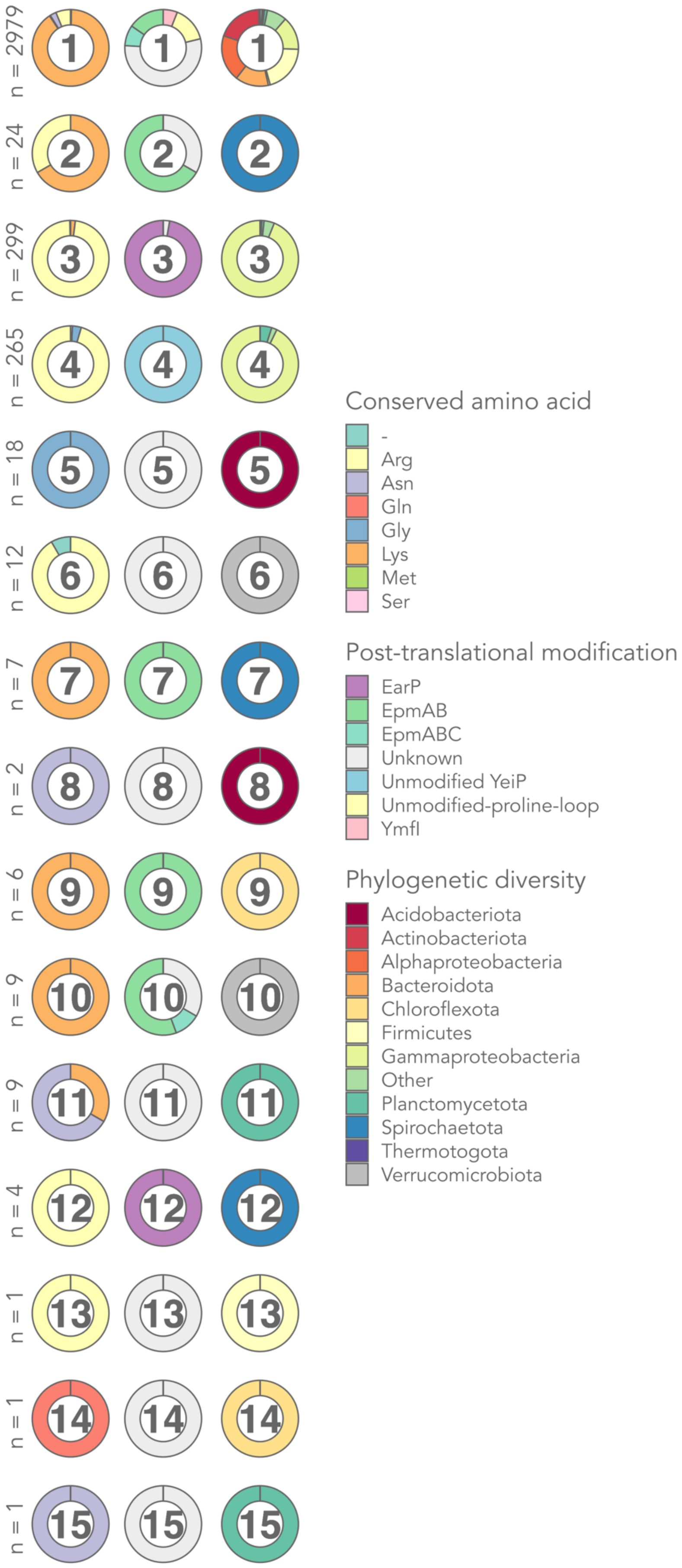


**Figure S1: Sequence homology clustering reveals many EFP types unique to specific phylogenetic clades.** To further characterize the large proportion of 'unknown' EFP types, we sorted all EFP sequences (3638 proteins) into 15 'families' using similarity network-based sequence homology clustering (*Methods*). We assigned protein sequences with ≥49% sequence identity and ≥80% sequence length alignment to the same family. While the majority of EFP fall into the first EFP family (1), including all EFP which are post-translationally modified at a lysine, there are many EFP families specific to certain phylogenetic clades. Each EFP family is represented three times and labelled with its family number. The three different columns of the figure report from left to right the conserved amino acid, the predicted post-translational modification, and phylogenetic placement of each EFP sequence, respectively. The number of proteins assigned to each family is indicated with "n = X".

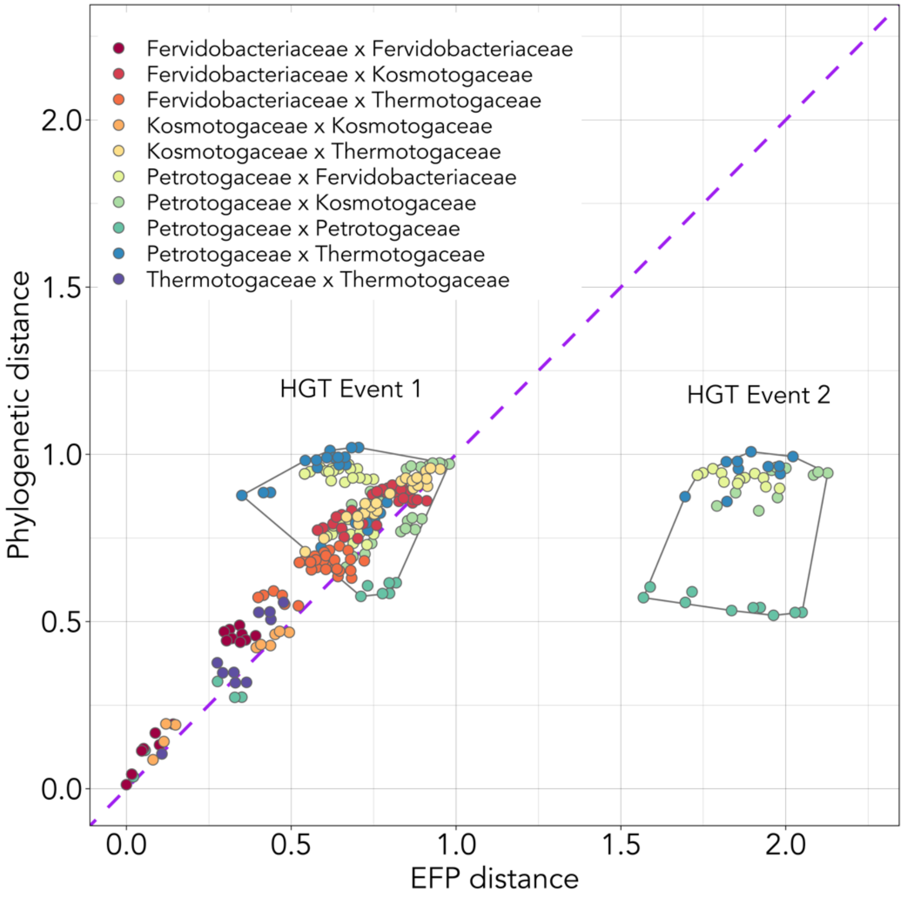


**Figure S2: Pairwise cophenetic distances between species' positions on the species and EFP protein sequence tree support horizontal transfer of EFP in the Thermotogota.**

The vertical axis shows the cumulative branch length between two species on the species tree (**Figure 2**), while the horizontal axis shows the cumulative branch length between the same two species on the EFP protein sequence tree (**Figure 3**). These phylogenetic distances correspond roughly to the number of amino acid substitutions per amino acid site. The color of the circles indicates from which phylogenetic families the pair of species being compared come from. For example, "Petrotogaceae x Thermotogaceae" indicates that a species from the family Petrotogaceae and a species from the family Thermotogaceae are being compared. We use this comparison as a visualization tool to identify potential horizontally transferred genes, as indicated by a phylogenetic distance that is not representative of the overall phylogenetic history of the clade. Two regions with anomalous evolutionary trajectories are outlined with dashed lines. They indicate EFP protein sequences that are more closely related than the background phylogeny (HGT event 1), and EFP protein sequences more distantly related than the background phylogeny of the species in which they are found (HGT event 2). Within the Petrotogaceae family two separate gene transfer events have occurred, one from within the family Thermotogaceae to the genera Petrotoga and Defluviitoga (HGT Event 1, Petrotogaceae x Thermotogaceae), and one from outside the phylum to the Oceanotoga and Geotoga genera (HGT Event 2, all colors).

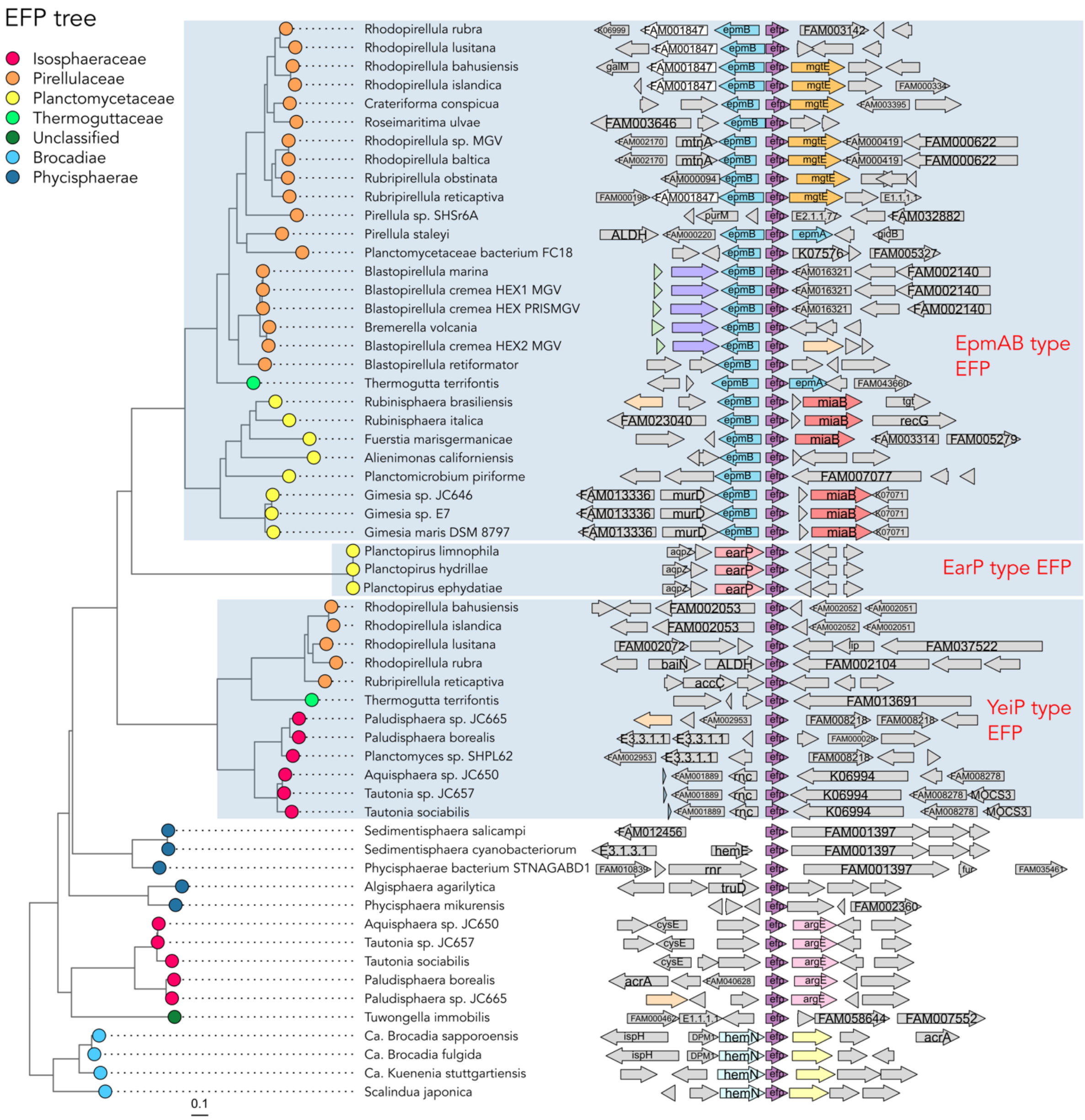


**Figure S3: Horizontal gene transfer of EFP in the Planctomycetota phylum. Left:** Phylogenetic tree of EFP amino acid sequences in the phylum Planctomycetota (*Methods*). Comparing this EFP tree with the phylogenetic tree in **Figure 4** shows that the phylogeny of EFP in the families Pirellulaceae, Thermoguttaceae, and Planctomycetaceae is not congruent with the overall phylogenetic tree of the phylum. Within the Planctomycetaceae family three separate gene transfer events of the *efp* gene have occurred (highlighted with grey-blue shading), one EarP type (middle box) and one EpmAB type (top box). Additionally, many genomes within the phylum encode a copy of the YeiP type EFP (bottom box). Some genomes encode two copies of EFP. **Right:** Gene synteny of each EFP gene from the tree on the left. For clarity, only select genes of interest and genes which occurred at least three times across all syntenies are indicated with color. Synteny of EFP (purple, centered) is not as well conserved among the Planctomycetota as in the Thermotogota. As in the Thermotogota, the EarP type EFP is co-located with its modifying enzyme (EarP, salmon). EpmB is co-located with its EFP type (EpmB, sky blue). EpmA (EpmA, sky blue) is co-located in only two species (*Pirellula staleyi* and *Thermogutta terrifontis*).

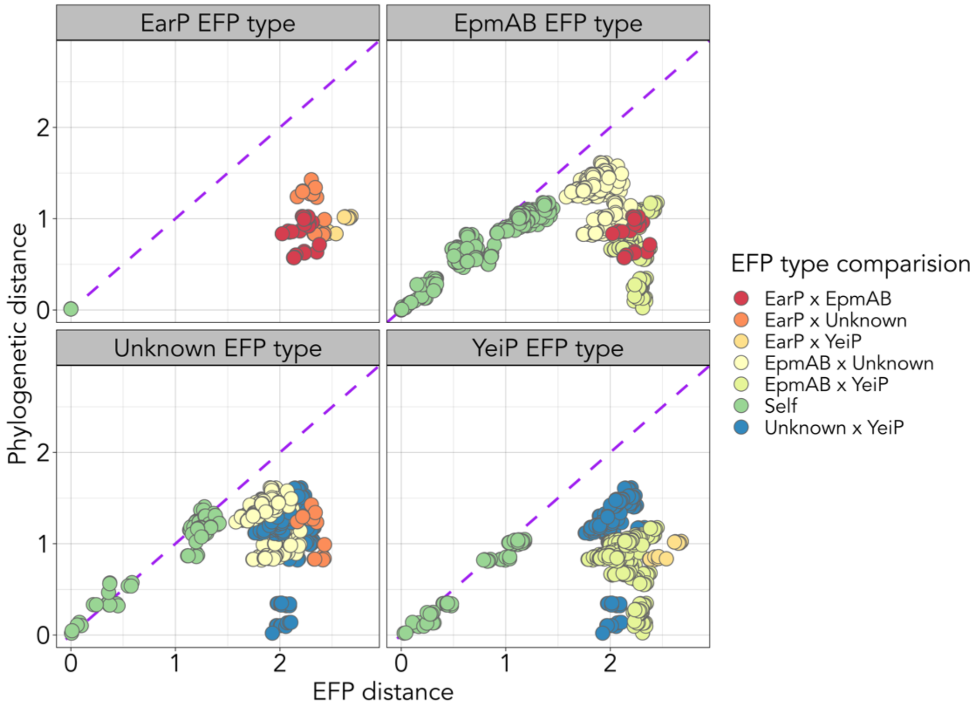


**Figure S4: Pairwise cophenetic distances between species' positions on the species and EFP protein sequence tree support horizontal transfer of EFP, as well as four distinct EFP types (EarP, EpmAB, YeiP, and Unknown type), in the Planctomycetota.** The vertical axis shows the cumulative branch length between two species on the species tree (**Figure 4**), while the horizontal axis shows the cumulative branch length between the same two species on the EFP protein sequence tree (**Figure S3**). These phylogenetic distances correspond roughly to the number of amino acid substitutions per amino acid site. The color of the circles indicates the EFP types being compared. For example, "EarP x EpmAB" indicates that an EarP type EFP and an EpmAB type EFP are being compared, while "Self" indicates that the same EFP types (EarP type x EarP type, for example) are being compared. We use this comparison as a visualization tool to identify potential horizontally transferred genes, as indicated by a phylogenetic distance that is not representative of the overall phylogenetic history of the clade. While within the same EFP type, the phylogenetic distances between EFP pairs tend to agree with the overall phylogenetic background (suggesting a single transfer event for each type), the sequence divergence between different EFP types is large.

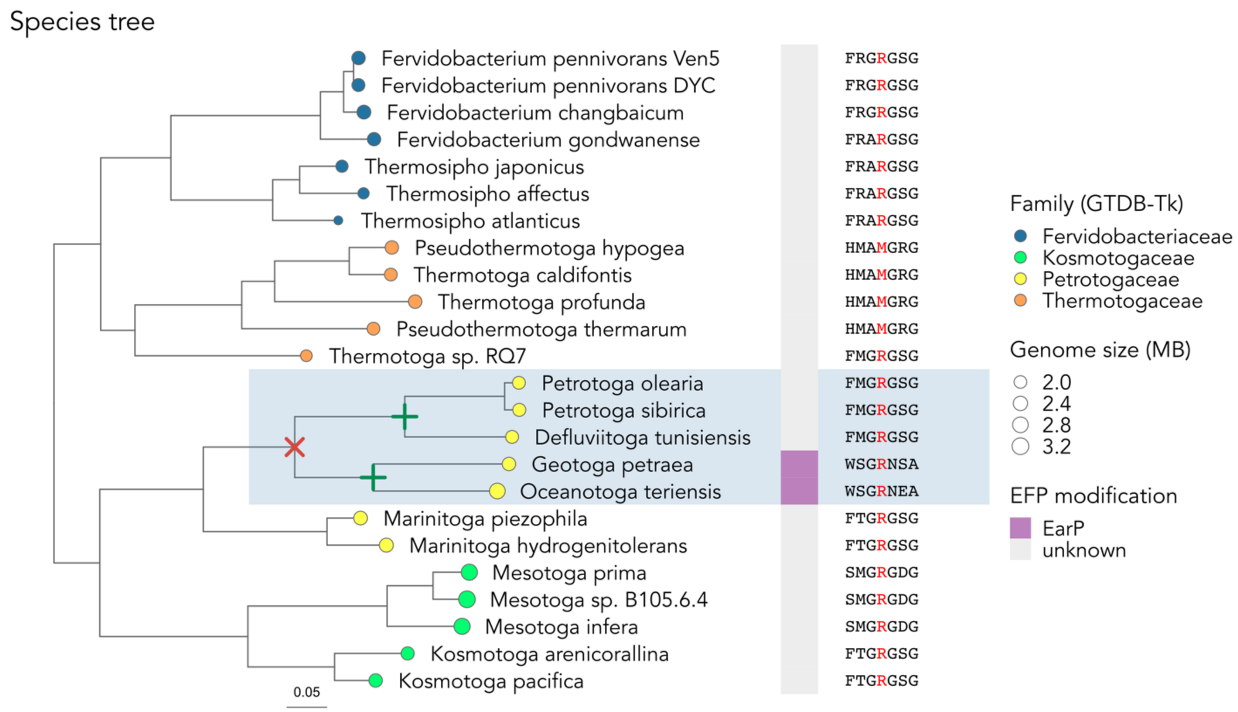


**Figure S5: Most native Thermotogota EFPs encode an arginine at the location of post-translational modification.** The position where the post-translational modification occurs is conserved in all known EFP. In most Thermotogota which do not show signs of EFP transfer, this residue is an arginine. We built this phylogenetic tree using amino acid sequences of 43 concatenated and conserved marker genes generated by CheckM (details in *Methods*). The color of the circle at the tree's tips represents the family these genomes belong to, according to the Genome Taxonomy Database (GTDB), as classified by the GTDB-Toolkit (GTDB-Tk). The size of the circle corresponds to relative genome size. Loss of 'native' EFP is indicated with a red X on the phylogenetic tree, while gain of a horizontally transferred EFP is indicated with a green +. The exact timing and order of these events is unknown. Species encoding horizontally transferred EFP are highlighted with grey-blue shading across the figure. The heatmap column to the right of the phylogenetic tree shows the bioinformatically identified EFP type. The sequence of the conserved loop region in domain I of EFP, which bears post-translational modifications if present, is displayed for the EFP of each species to the right of the heatmap (equivalent to *E. coli* EpmAB type EFP positions K31:A37).

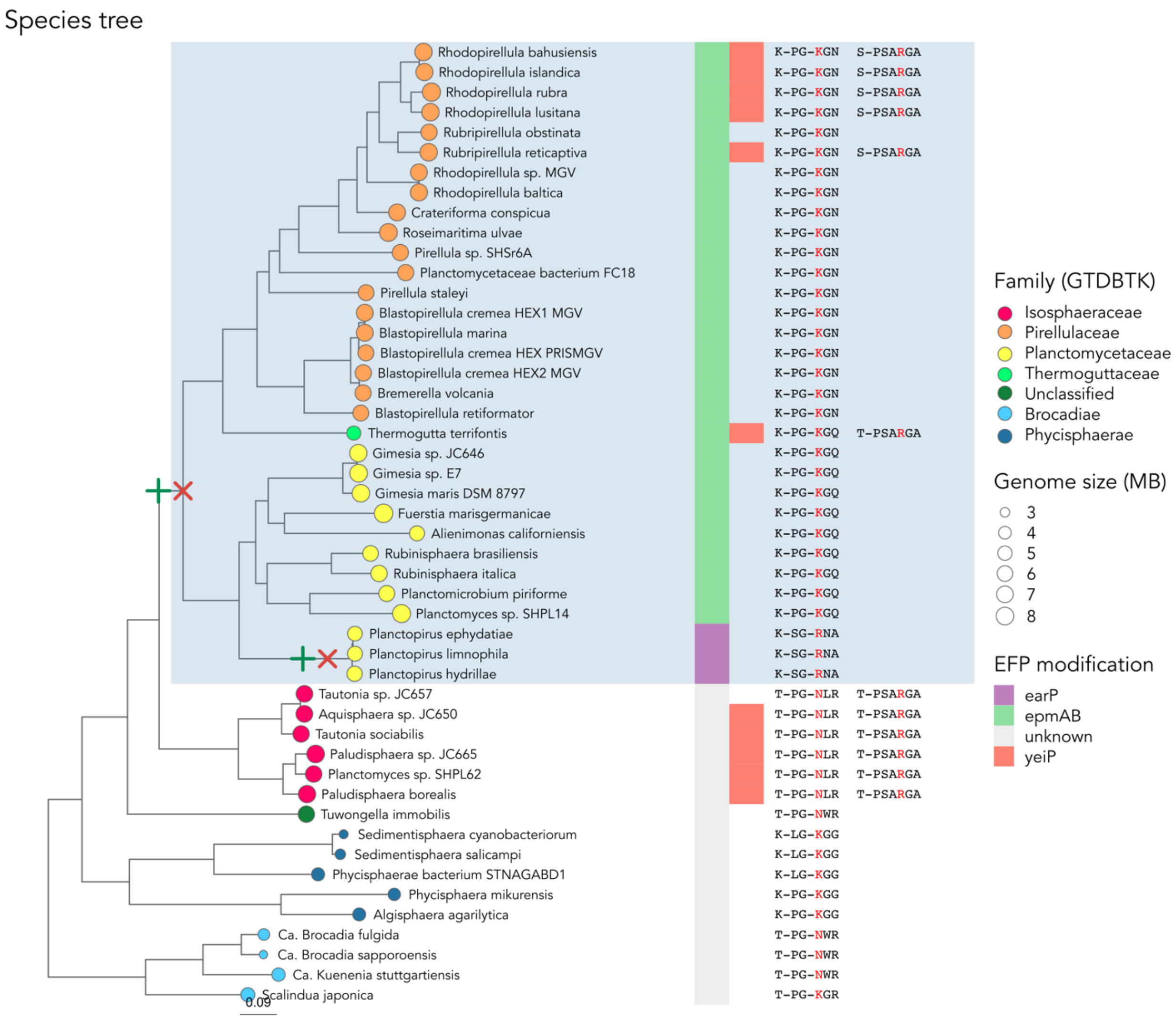


**Figure S6: Native Planctomycetota EFP encode lysine or asparagine at the location of post-translational modification.** The position of the amino acid where the post-translational modification of EFP is attached is conserved in all known EFP. In Planctomycetota species which do not show obvious signs of EFP transfer, this residue is either an asparagine or a lysine. We built this phylogenetic tree using amino acid sequences of 43 concatenated and conserved marker genes generated by CheckM[1] (*Methods*). The color of the circle at the tree endpoints represents the family these genomes belong to according to GTDBTK and the size of the circle corresponds to relative genome size. Loss of native EFP is indicated with a red X on the phylogenetic tree, while gain of a horizontally transferred EFP is indicated with a green +. The exact timing and order of these events is unknown. HGT events of the *efp* gene are highlighted with blue-grey shading across the figure. The first column of the heatmap on the right shows the bioinformatically identified EFP types. YeiP is present as a secondary EFP in multiple genomes. The sequence of the conserved loop region in domain I of EFP, which bears post-translational modifications if present, is displayed in the final two columns with order corresponding to the heatmap (sequence is equivalent to *E. coli* EpmAB type EFP positions K31:Q36).

**Supplemental Table S1:** Level of conservation of polyproline motifs found to be sensitive to EFP transfer (**Figures 3 & 4**) in our wider 3000 genomes dataset. Not every genome contains each protein, and some genomes contain multiple copies of some proteins.

| Protein | KEGG KO | Fraction of proteins containing conserved PP motif |
|---|---|---|
| ValS | K01873 | 99.8 % (2902/2907) |
| IleS1 | K01870 | 99.7 % (1958/1964) |
| IleS2 | K01870 | 99.3 % (1343/1353) |
| Lon | K01338 | 98.8 % (3100/3139) |
| ClpC | K03696 | 97.9 % (2197/2244) |
| FtsH | K03798 | 85.5 % (3666/4287) |
| TrpB | K06001 | 50.0 % (262/524) |
| RpoD | K03086 | 5.7 % (248/4344) |
| YcaJ | K07478 | 3.9 % (123/3177) |
| PilC | K02653 | 1.2 % (18/1477) |

### Supplemental results

#### Sequence homology clustering reveals EFP families unique to specific phylogenetic clades

We found three EFP families specific to the Spirochaetota (FAM02, containing 16 lysine-EpmAB type EFPs, and 8 arginine type EFPs with an unknown modification system; FAM07 containing 7 lysine-EpmAB type EFPs; and FAM12 containing 4 sequences of the arginine-EarP type, **Figure S1**). In addition, we found several unique EFP families of completely unknown modification type. For example, FAM05 includes 18 Acidobacteria glycine type EFPs; FAM06 includes 12 Verrucomicrobiota arginine type EFPs, FAM08 includes 2 asparagine type EFPs unique to the family Holophagaceae from the phylum Acidobacteria, and FAM11 includes 9 Planctomycetota lysine/asparagine type EFP. Moreover, we found two small clades that co-occur with EpmAB modification systems and feature a lysine as their conserved residue: FAM09 contains 6 Chloflexota EFPs and FAM10 contains 9 EFPs specific to the class Chlamydiia. Lastly, there were three species with unique EFPs: the Planctomycete *Tuwongella immobilis*, the Chloroflexi *Thermoflexus hugenholtzii*, and the Mycoplasma Candidatus *Hepatoplasma crinochetorum*. These sequences may have been clustered independently due to poor sequence representation because their genomes were the sole representatives from their families in our dataset.

We also found several cases that appear to be reversions of EFP from modified forms to unmodified forms. For example, FAM02 of the Spirochaetota contains a majority of lysine-EpmAB type EFPs, but also contains some arginine type EFPs. These EFPs are exclusive to the Borreliaceae, a family whose genomes contain many pseudogenes and undergo recurrent recombination with extrachromosomal elements[1]. It may be that one gene of the modification system was lost, leading to mutation of EFP when selection for the lysine residue was relieved.

#### EFP pairings

After the lysine-EpmAB type EFP and arginine-YeiP type EFP, the next most common EFP pairing is two distinct EFP types of unknown modification type, (49 genomes, most common among Acidobacteriota). We found some cases of lysine-EpmAB type EFPs co-occurring with unknown EFP types (43 genomes, most common among the Alphaproteobacteria with 25 genomes). These unknown EFP types often encode a lysine at the conserved position and a neighboring proline (P32 in *E. coli*[2]) known to be important for the EpmAB type modification (26 cases, in the Alphaproteobacteria orders Rhizobiales and Rhodobacterales, and the myxobacteria *Myxococcus fulvus*). This raises the possibility of two lysine-EpmAB type EFP coexisting in the same genome. Rare pairings in our dataset include an arginine-YeiP type EFP with an unknown EFP type (8 genomes, all Planctomycetota), and an arginine-EarP type EFP co-existing with a lysine EpmAB type (4 genomes, all Gammaproteobacteria).


1. Casjens, S. *et al.* A bacterial genome in flux: The twelve linear and nine circular extrachromosomal DNAs in an infectious isolate of the Lyme disease spirochete Borrelia burgdorferi. *Mol Microbiol* **35**, 490–516 (2000).
2. Volkwein, W. *et al.* Switching the post-translational modification of translation elongation factor EF-P. *Front Microbiol* **10**, (2019).
